# *P16* methylation increases the sensitivity of cancer cells to the CDK4/6 inhibitor palbociclib

**DOI:** 10.1101/771337

**Authors:** Paiyun Li, Xuehong Zhang, Liankun Gu, Jing Zhou, Dajun Deng

## Abstract

The *P16* (*CDKN2A^ink4a^*) gene is an endogenous CDK4/6 inhibitor. Palbociclib (PD0332991) is an anti-CDK4/6 chemical for cancer treatment. *P16* is most frequently inactivated by copy number deletion and DNA methylation in cancers. It is well known that cancer cells with *P16* deletion are more sensitive to palbociclib than those without. However, whether *P16* methylation is related to palbociclib sensitivity is not known. By analyzing public pharmacogenomic datasets, we found that the IC50 of palbociclib in cancer cell lines (n=522) was positively correlated with both the *P16* expression level and *P16* gene copy number. Our experimental results further showed that cancer cell lines with *P16* methylation were more sensitive to palbociclib than those without. To determine whether *P16* methylation directly increased the sensitivity of cancer cells to palbociclib, we induced *P16* methylation in the lung cancer cell lines H661 and HCC827 and the gastric cancer cell line BGC823 via an engineered *P16*-specific DNA methyltransferase (P16-Dnmt) and found that the sensitivity of these cells to palbociclib was significantly increased. The survival rate of P16-Dnmt cells was significantly lower than that of vector control cells 48 hrs post treatment with palbociclib (10 μM). Notably, palbociclib treatment also selectively inhibited the proliferation of the *P16*-methylated subpopulation of P16-Dnmt cells, further indicating that *P16* methylation can increase the sensitivity of cells to this CDK4/6 inhibitor. These results were confirmed in an animal experiment. In conclusion, inactivation of the *P16* gene by DNA methylation can increase the sensitivity of cancer cells to palbociclib.

## Introduction

The ability to sustain uncontrolled cell proliferation is one of the hallmarks of cancer cells [1]. The normal process of cell division depends on the cell cycle, a series of highly regulated steps manipulated by a set of specific cyclins that act in association with cyclin-dependent kinases (CDKs) [2–4]. The CDK4/6 complex plays a key role in cell cycle progression via monophosphorylation of retinoblastoma protein (RB) and subsequently promotes G1-S phase transition [5, 6].

The clinical implementation of first-generation nonselective CDK inhibitors was originally hampered by the high toxicity and low efficacy of these agents [7, 8]. Second-generation selective CDK4/6 inhibitors, including palbociclib, ribociclib, and abemaciclib, can induce reversible G1 phase cell cycle arrest in RB-positive tumor models with improved effectiveness and reduced adverse effects [9–17]. On the basis of the significant improvements in progression-free survival (PFS) in the PALOMA-1, MONALEESA-2 and MONARCH-1 and 2 clinical trials, palbociclib, ribociclib, and abemaciclib received FDA approval for the treatment of hormone receptor (HR)-positive and human epidermal growth factor receptor 2 (HER2)-negative breast cancer [18–20]. However, not all of these patients could benefit from treatment with CDK4/6 inhibitors [21, 22]. Therefore, biomarkers for predicting the response to these drugs are needed.

The P16 protein, encoded by the *CDKN2A^ink4a^* gene, is an endogenous cellular CDK4/6 inhibitor that controls the G1-S phase transition of the cell cycle. This gene is one of the most frequently inactivated genes in cancer genomes; it is inactivated mainly by DNA methylation [23]. It has been reported that cancer cells with *P16* copy number deletion are more sensitive to palbociclib than those without [24–28]. Hence, we sought to determine whether cancer cells with *P16* methylation exhibit increased sensitivity to therapeutic CDK4/6 inhibitors. Building on these premises, we systematically investigated the relationship between *P16* methylation and the sensitivity of cancer cells to the CDK4/6 inhibitor palbociclib using both public datasets and cell models of *P16* methylation induced by an artificial *P16*-specific methyltransferase (P16-Dnmt) [29].

## Materials and methods

### Dataset sources

*P16* (*CDKN2A^ink4a^*) gene expression levels and copy numbers in cancer cell lines were downloaded from the Cancer Cell Line Encyclopedia (CCLE) [https://portals.broadinstitute.org/ccle] [30]. The half-maximal inhibitory concentration (IC50) value data for the CDK4/6 inhibitor palbociclib (PD0332991) in various cell lines were downloaded from the Genomics of Drug Sensitivity in Cancer (GDSC) database [https://www.cancerrxgene.org/translation/Drug/1054#t_IC50] [31].

### Cell lines and culture

Human lung cancer cell lines H1975, H1395, H661, H596, H460, H358, H292 and HCC827, as well as human breast cancer cell lines HCC1937, MDA-MD-468 and MDA-MD-157, were purchased from the National Infrastructure of Cell Line Resource (Institute of Basic Medical Sciences, Chinese Academy of Medical Sciences). Human lung cancer cell lines H1299 and A549; human breast cancer cell lines MDA-MB-231 and MCF7; human gastric cancer cell lines BGC823, MGC803 and AGS; human colon cancer cell lines HCT116, RKO and SW480; and human liver cancer cell line HepG2 were obtained from laboratories at Peking University Cancer Hospital and Institute. The H596 and MCF7 cell lines were cultured in DMEM (Gibco, USA) containing 10% FBS (Gibco) and 1% penicillin-streptomycin (Gibco), the AGS cell line was cultured in F12 medium (Gibco), and the remaining cell lines were cultured in RPMI 1640 medium (Gibco). All cells were maintained at 37 °C in humidified air with 5% CO_2_.

### Cell viability assay

The CDK4/6 inhibitor palbociclib (PD0332991) was purchased from Selleck Chemicals (#S1116; USA) and dissolved in double-distilled H2O as a stock solution at concentration of 10 mM. Cells were seeded in 96-well plates at 4,000 cells per well (4 wells/group). Forty-eight hrs post treatment with various concentrations of palbociclib, cell confluence was evaluated and analyzed using the IncuCyte ZOOM system (Essen BioSci, USA). The half-maximal inhibitory concentration (IC50) was calculated using GraphPad Prism 6 software.

### Transfection of P16-Dnmt and empty control vector

The established P16-Dnmt expression vector and pTRIPZ empty control vector were prepared and used to transfect cells as previously described [29]. Briefly, to induce methylation of CpG islands around the *P16* transcription start site, an engineered *P16* promoter-specific seven zinc finger protein (7ZFP) was fused with the catalytic domain of mouse Dnmt3a (approximately 608–908 aa) and integrated into the pTRIPZ vector, which contained a “Tet-On” switch (Open Biosystem, USA). The lung cancer cell lines H661 and HCC827 and the gastric cancer cell line BGC823 were infected with lentiviral particles containing the P16-Dnmt or control vector and incubated for 48 hrs. Then, puromycin (Sigma, USA) was added to the medium (final concentration, 1 μg/mL) to kill nontransfected cells. The pooled cells treated with puromycin for two weeks were considered stably transfected cells. Then, these cells were treated with 0.25 μg/mL doxycycline (Sigma, USA) for 14 days to induce P16-Dnmt expression.

### DNA extraction and bisulfite modification

Genomic DNA was extracted from cells or tumor tissues and subjected to bisulfite treatment using an EZ DNA Methylation-Gold Kit (ZYMO RESEARCH, USA) according to the manufacturer’s instructions. The modified DNA was stored at −20 °C until use.

### Methylation-specific PCR (MSP) and MethyLight assay

The methylation status of *P16* CpG islands was assessed by a 150/151-bp methylation-specific PCR (MSP) or 115-bp quantitative MethyLight assay as previously described [32–34]. Briefly, bisulfite-modified genomic DNA was amplified using the methylated/unmethylated *P16*-specific primer sets in the MSP assay or using the *P16*-specific primers and probe in the 115-bp MethyLight assay (Table 1). All PCR products were amplified with HotStart Taq DNA polymerase (QIAGEN, Germany). In the MethyLight assay, the level of *P16* methylation was determined using an Applied Biosystems 7500 Real-Time PCR System (Applied Biosystems, USA) and normalized to that of the internal control gene, *COL2A1* (*collagen type II alpha 1*). The relative copy number (RCN) of the methylated *P16* CpG islands was calculated according to the formula [2^−ΔCt^, (ΔCt=Ct_methylated-*P16*_ – Ct_COL2A1_)].

**Table 1.**
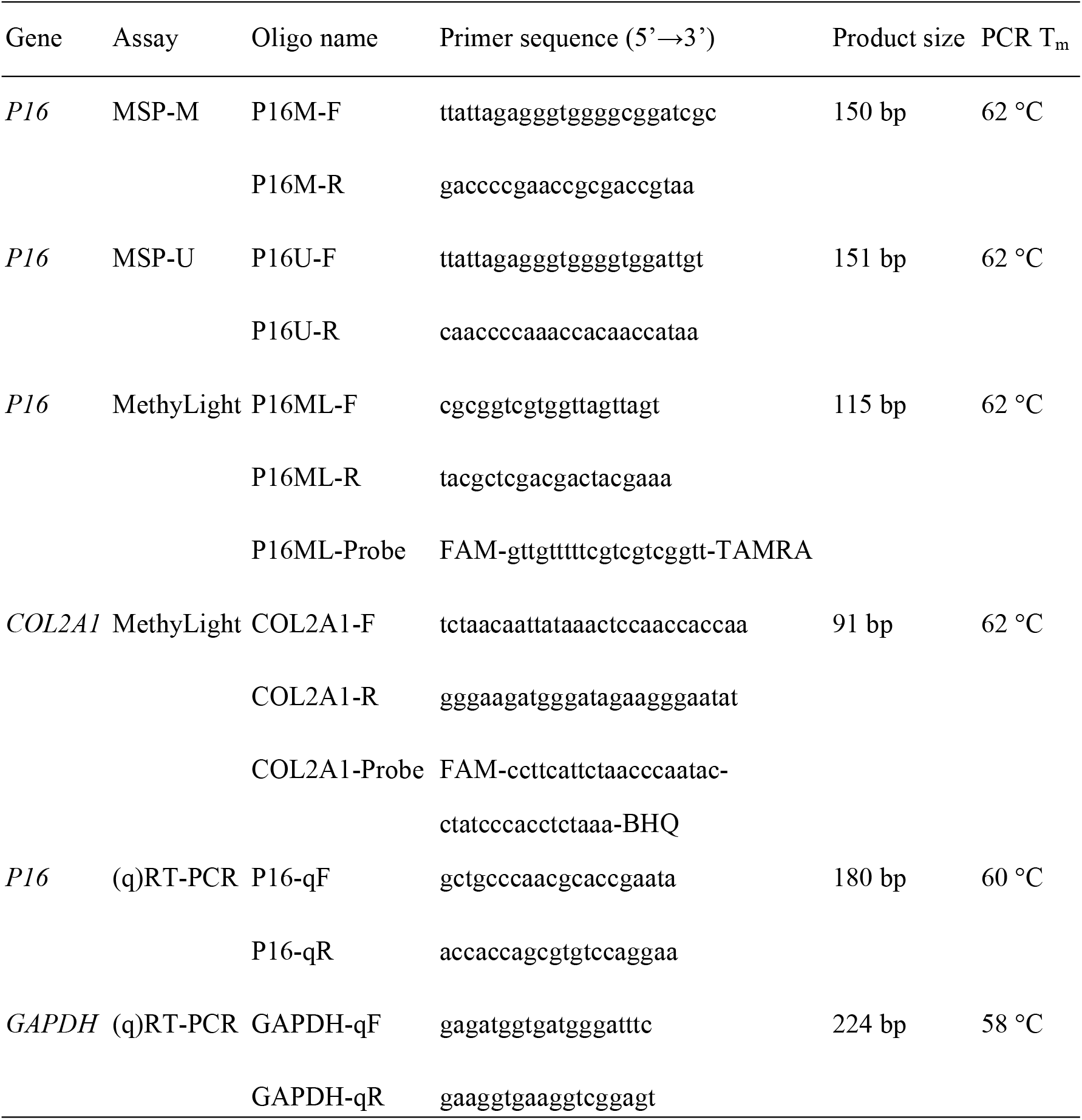
Sequences of oligonucleotides used as primers and probes in various PCR assays

### RNA extraction and quantitative RT-PCR (qRT-PCR)

Cells were harvested at a confluence of approximately 70%. Total RNA was extracted from transfected cells or tumor tissues by TRIzol reagent (Thermo Fisher, USA) and reverse transcribed using TransScript First-Strand cDNA Synthesis SuperMix (TransGen, China). The qRT-PCR for *P16* mRNA was performed using primer sets shown in Table 1. *GAPDH* was used as the reference gene. The amplification was performed with FastStart Universal SYBR Green Master (ROX) (Roche, Switzerland) in the Applied Biosystems 7500 Real-Time PCR System, as previously described [35].

### Immunofluorescence and confocal microscopy analysis

Cells were fixed in 4% polyformaldehyde for 10 min at room temperature, treated with 1% Triton X-100 in PBS for 10 min, blocked with 5% bovine serum albumin (BSA) for 1 hr, and hybridized to a mouse monoclonal antibody against the P16 protein (Ventana Roche Diagnostics, E6H4, Switzerland) overnight at 4°C. Samples were incubated with FITC-labeled secondary antibody (KPL, 172-1806, USA) for 1 hr at room temperature, followed by nuclear staining with DAPI. Fluorescence images were acquired and analyzed with an ImageXpress Micro High Content Screening System (Molecular Devices, USA).

### Western blot analysis

Primary monoclonal antibodies against RB (Abcam, ab181616, UK), Ser780-phosphorylated RB (p-RB) (Cell Signaling Technology, #9307, USA), P16 (Abcam, ab108349), and GAPDH (Proteintech Group, 60004-1-Ig, USA) were diluted at 1:1000, 1:1000, 1:1000 and 1:15000, respectively. Signals were visualized using Immobilon Western HRP Substrate (Millipore, USA).

### BALB/c nude mouse xenograft model and treatment

HCC827 cells (2 × 10^6^ cells in 200 μL of PBS) stably transfected with the P16-Dnmt vector or control vector and induced with 0.25 μg/mL doxycycline for 7 days were injected subcutaneously into the lower limbs of BALB/c nude mice (female, 5 weeks old, weighing 18-22 g, purchased from Beijing Huafukang Biotech). Mice were provided access to distilled, sterile water containing 2 μg/mL doxycycline ad libitum. When the tumor size reached approximately 200 mm^3^, on the 30th day post transplantation, mice were randomized into the control group and the palbociclib group (6 mice/group) and treated with control buffer or palbociclib [100 mg/kg in 50 mM sodium lactate buffer (pH 4.0)] via oral gavage (i.g.) daily for 3 weeks. Then, mice were sacrificed, and the tumors were weighed. Tumor samples were processed for formalin-fixed and paraffin-embedded (FFPE) sections or Western blot analysis.

### Immunohistochemistry (IHC)

For IHC, after dewaxing, rehydration, endogenous peroxidase quenching and blocking with 5% BSA according to standard procedures, 4-μm thick formalin-fixed, paraffin-embedded (FFPE) sections were incubated with primary antibody against Ki-67 (1:300) overnight at 4 °C. The remaining IHC steps were the same as previously described [36].

### Statistical analysis

Statistical analyses were performed using SPSS 23 software or GraphPad Prism 6 software. The Kolmogorov–Smirnov test and Shapiro–Wilk test were used to estimate the normality of distributions. The relationship between the IC50 value of palbociclib and the relative *P16* (*CDKN2A^ink4a^*) mRNA level or copy number in various cancer cell lines was assessed using the nonparametric Spearman correlation test and a linear regression model. The Mann–Whitney test was used for nonnormally distributed data. Student’s t-test was used for normally distributed data. Statistical significance was accepted at *p*<0.05.

## Results

### P16 methylation is positively correlated with high sensitivity of cancer cells to palbociclib

To explore the relationship between *P16* methylation and the sensitivity of cancer cells to palbociclib, we first analyzed the correlation between the IC50 levels of palbociclib in 522 cell lines and the transcription level and copy number of the *CDKN2A/P16* gene using public pharmacogenomics datasets [30, 31]. The analysis results revealed that the palbociclib IC50 levels in these cell lines were positively correlated with the *CDKN2A* mRNA level (*r*=0.3133, *p*<0.0001; Fig 1a) and *CDKN2A* copy number (*r*=0.2832, *p*<0.0001; Fig 1b). This result suggests that cancer cell lines with low or no *P16* expression are more sensitive to palbociclib than those with high *P16* expression.

**Fig 1.**
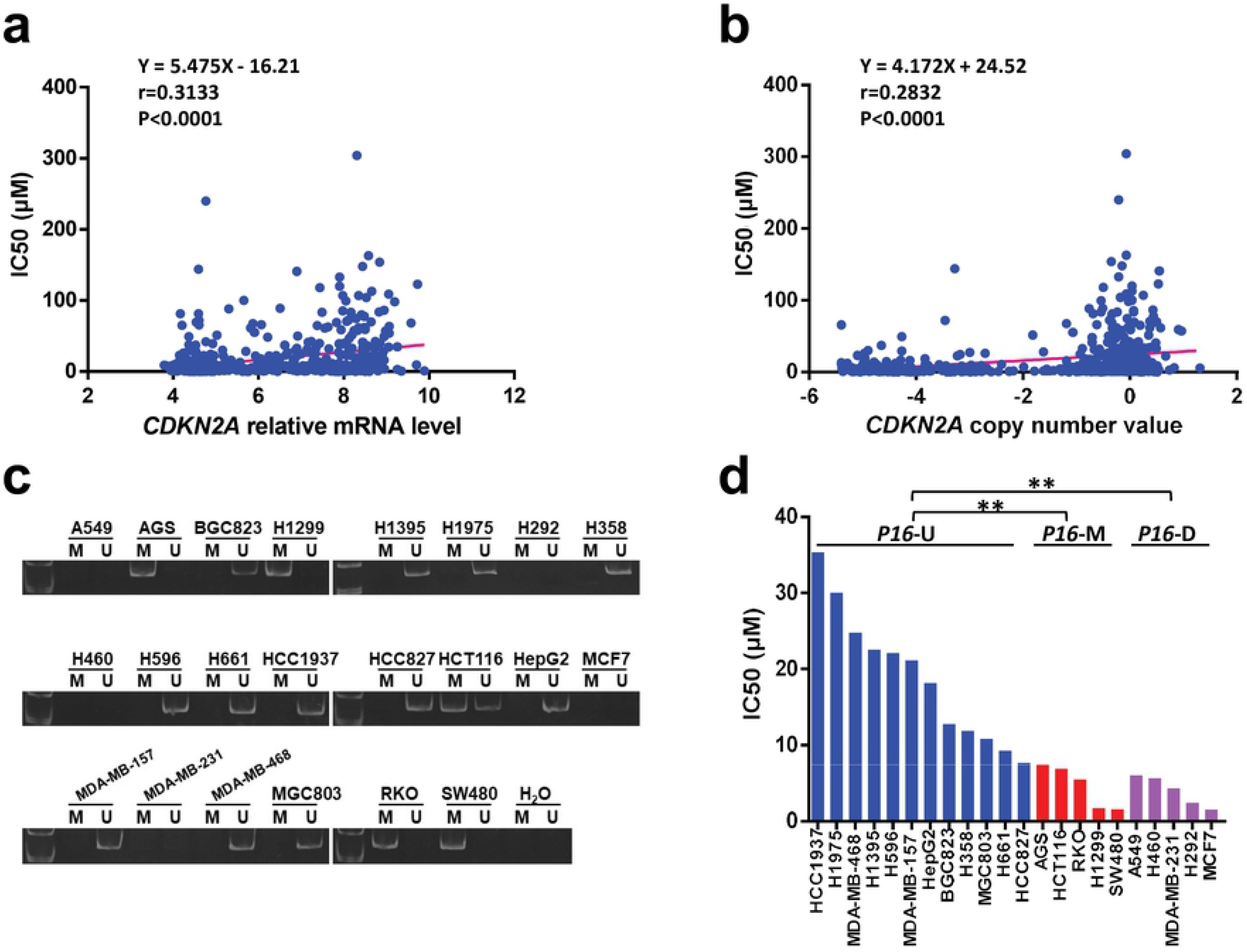
Association between *P16* expression and methylation and palbociclib sensitivity in cancer cell lines. (**a** and **b**) Correlations between the palbociclib IC50 levels and *CDKN2A/P16* mRNA levels and copy number in cancer cell lines (*n*=522). The *p*-values were measured by a nonparametric Spearman correlation test and linear regression model. (**c**) The methylation status of the *P16* gene in different cell lines as assessed by MSP. (**d**) The palbociclib IC50 values in 22 cancer cell lines with different *P16* alterations, including *P16* copy number deletion (*P16*-D), *P16* methylation (*P16*-M), and *P16* unmethylated and undeleted (*16*-U). Student’s t-test, \*\**p*<0.01.

Then, we validated the correlation in 22 cancer cell lines, which were divided into *P16*-unmethylated (*P16*-U), *P16*-methylated (*P16*-M), and *P16*-deleted (*P16*-D) groups according to the results of MSP (Fig 1c). We found that the IC50 values in the *P16*-U cell lines (n=12) were consistently higher than those in the *P16*-M cell lines (n=5) and *P16*-D cell lines (n=5) [18.88 *vs*. 4.61 and 3.99 (μM), *p*<0.01; Fig 1d]. There was no difference in the IC50 values between the *P16*-M and *P16*-D cell lines. These results indicate that similar to copy number deletion, *P16* methylation is positively associated with high sensitivity of cancer cells to palbociclib.

### P16-specific methylation directly increases the sensitivity of cancer cells to palbociclib

To determine whether *P16* methylation directly increases the sensitivity of cancer cells to palbociclib, we stably transfected three *P16*-U cell lines (including two lung cancer cell lines, H661 and HCC827, and one gastric cancer cell line, BGC823) with the P16-Dnmt vector and induced P16-Dnmt expression in these cells by doxycycline treatment for 2 weeks. *P16* methylation was successfully induced in these P16-Dnmt cells, as evidenced by MSP (Fig 2a). qRT-PCR analysis revealed that the level of *P16* expression was markedly decreased in these P16-Dnmt cells (Fig 2b). Then, these cells were treated with palbociclib at different concentrations (0, 5, 10 μM), and cell viability (survival rate) was assessed. The results showed that epigenetic inactivation of the *P16* gene by P16-Dnmt significantly decreased the viability of palbociclib-treated cells (Fig 2c). Forty-eight hrs post treatment with palbociclib (final concentration, 10 μM), the survival rate of P16-Dnmt cells was significantly decreased compared to that of vector control cells: 48.6% *vs*. 56.9% for H661 cells, 19.9% vs. 34.5% for HCC827 cells, and 52.5% vs. 68.3% for BGC823 cells. These results indicate that *P16*-specific methylation can directly increase the sensitivity of cancer cells to palbociclib.

**Fig 2.**
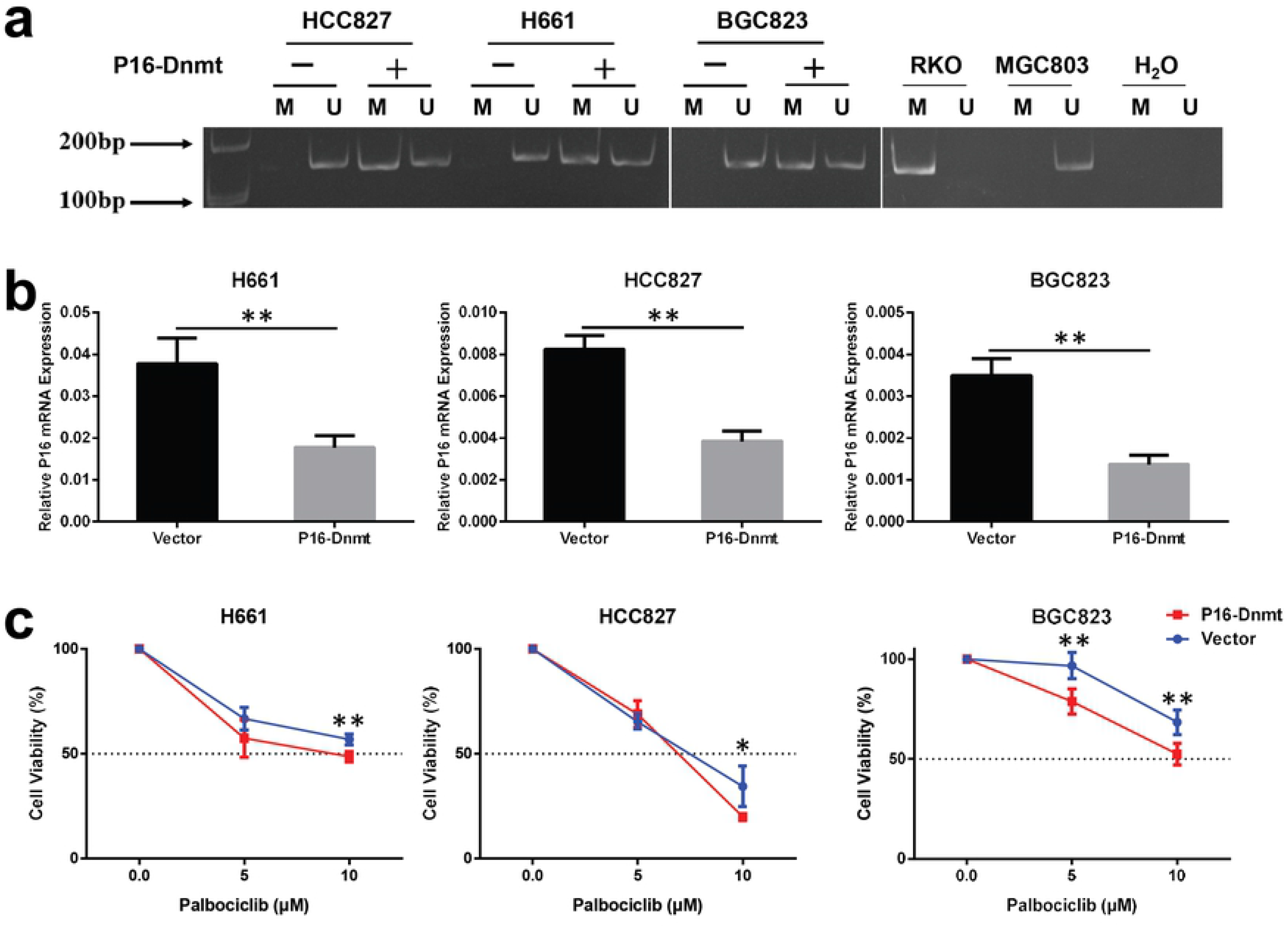
Effect of *P16*-specific methylation on the sensitivity of cancer cells to palbociclib. (**a**) Methylation of the CpG *P16* island in three cancer cell lines stably transfected with an engineered *P16*-specific DNA methyltransferase (P16-Dnmt) by MSP. Genomic DNA samples from RKO and MGC803 cells were used as the positive and negative controls for *P16* methylation, respectively. (**b**) Effect of *P16*-specific methylation on the levels of *P16* expression in cell lines stably transfected with P16-Dnmt, as assessed by qRT-PCR. (**c**) Effect of *P16*-specific methylation on the viability (survival rate) of cells treated with palbociclib for 48 hrs, as assessed with the IncuCyte ZOOM system. Each point represents the mean ± SD of 4 wells. Student’s t-test, \**p*<0.05, \*\**p*<0.01.

### Selective inhibition of the proliferation of P16-methylated cancer cells by palbociclib in vitro

As described above, *P16* inactivation by methylation was not detected in approximately one/third of P16-Dnmt cells (Fig 2b). Therefore, we further investigated whether palbociclib treatment could selectively inhibit the proliferation of the *P16*-methylated subpopulation of P16-Dnmt cells. As expected, the population of *P16*-methylated cells was significantly decreased by palbociclib treatment in the H661, HCC827, and BGC823 cell lines stably transfected with P16-Dnmt (Fig 3a). In contrast, the *P16* mRNA level was significantly increased (Fig 3b) by palbociclib treatment, while the *P16* mRNA level in the vector control cells was not affected. Furthermore, immunofluorescence and confocal microscopy analyses confirmed the qRT-PCR results. An increase in the number of P16 protein-positive cells was observed in these P16-Dnmt cell lines treated with palbociclib (Figs 3c, S1 and S2). Such a change was not observed in vector control cells. These results show that the proliferation of *P16*-methylated (or P16-negative) cells is selectively inhibited by palbociclib treatment.

**Fig 3.**
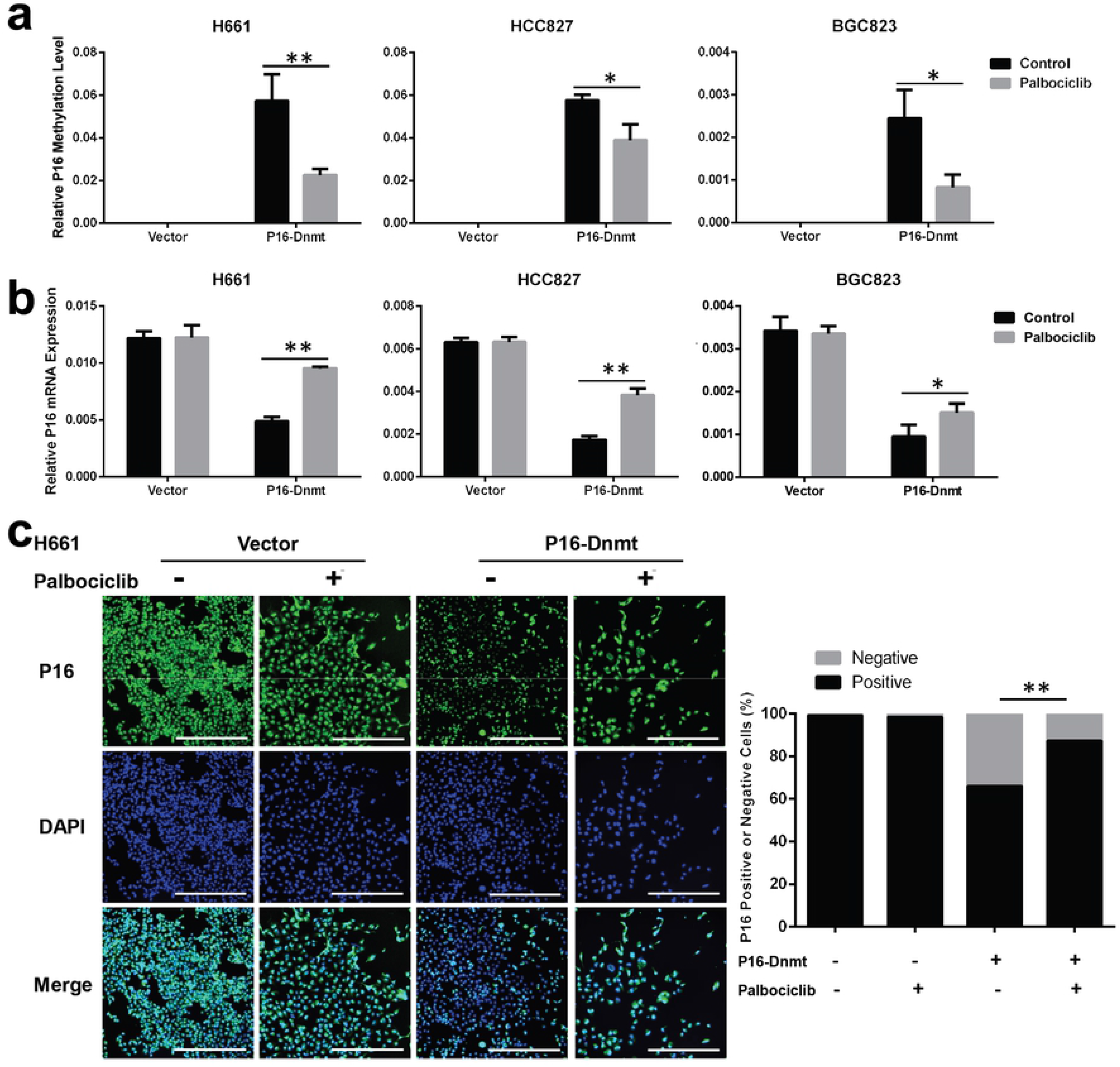
Selective inhibition of the proliferation of *P16*-methylated cancer cells by palbociclib *in vitro*. After palbociclib treatment (H661 and BGC823 cells: 10 μM; HCC827 cells: 7 μM) for 48 hrs, cells were harvested at 70–80% confluence. (**a** and **b**) Effect of palbociclib treatment on the levels of *P16*-methylated alleles and *P16* mRNA in cell lines stably transfected with P16-Dnmt, as assessed by MSP and qRT-PCR assays, respectively. The data are presented as the means ± SDs. (**c**) The results of immunofluorescence and confocal microscopy analyses to directly detect changes in the subpopulation of cells staining positive for P16 protein within the H661 cell line stably transfected with P16-Dnmt. Scale bar, 400 μm. Student’s t-test, \**p*<0.05, \*\**p*<0.01.

### Selective inhibition of the growth of P16-methylated cancer cells by palbociclib in nude mice

To verify the selective inhibition of *P16*-methylated cancer cells by palbociclib described above, HCC827 cells stably transfected with P16-Dnmt were subcutaneously transplanted into nude mice. When the xenograft tumor size reached approximately 200 mm^3^, mice were randomized and treated with palbociclib (100 mg/kg, i.g.) or control buffer for three weeks. Compared with control buffer, palbociclib significantly inhibited the growth of P16-Dnmt tumors (*p*<0.01; Fig 4a). Consistent with this result, the proportion of Ki67-positive cells was significantly decreased in P16-Dnmt tumors treated with palbociclib but not in control vector tumors (Fig 4b).

**Fig 4.**
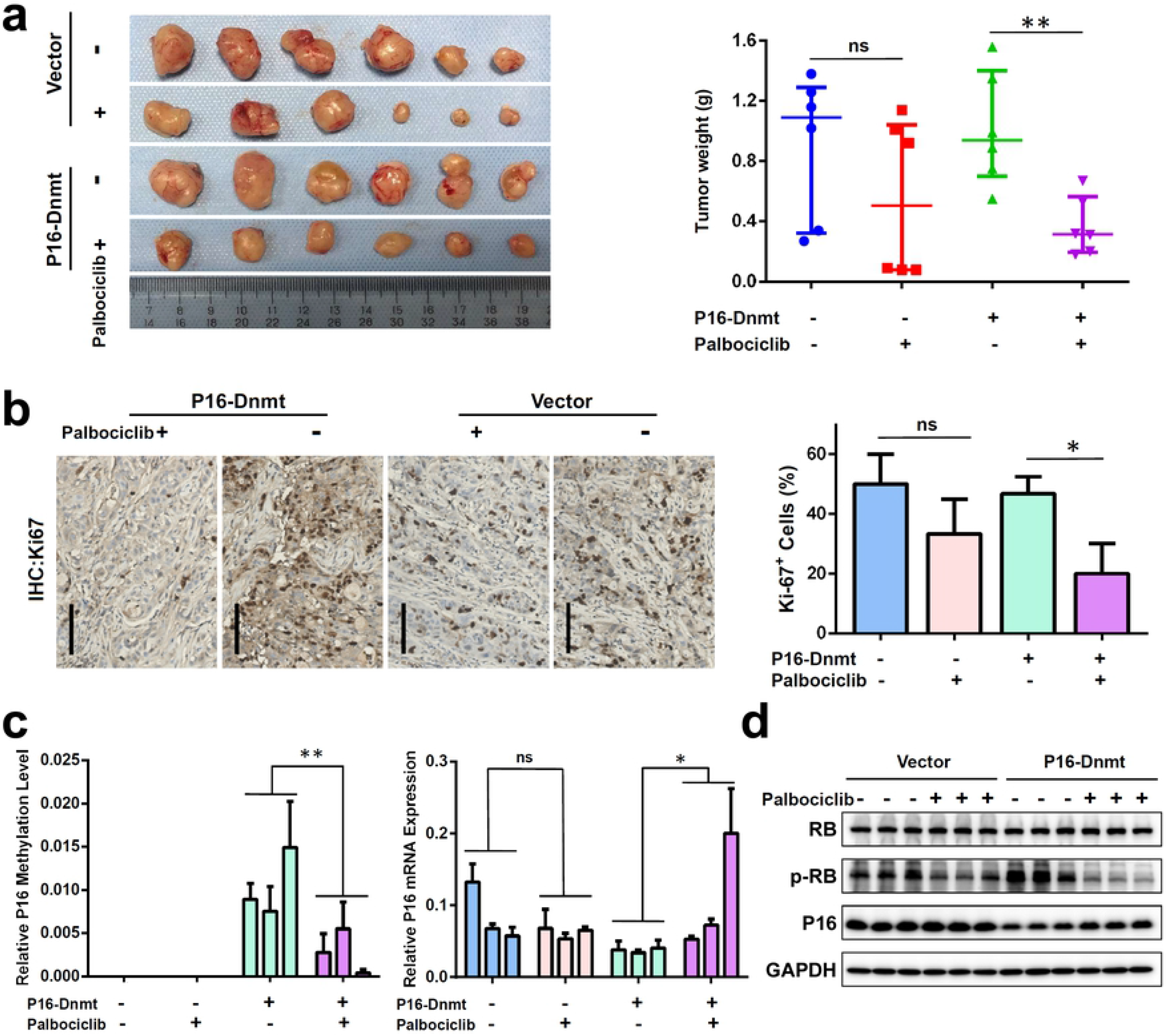
Effect of palbociclib treatment on the growth of tumors derived from HCC827 cancer cells with and without stable transfection of P16-Dnmt in nude mice. (**a**) Images of HCC827 xenograft tumors in different groups of mice (6 mice/group). These mice were treated with palbociclib (100 mg/kg, i.g.) or control buffer for 3 weeks. A tumor weight chart is inserted on the right side. The data are expressed as the medians ± interquartile ranges; Mann–Whitney test, \*\**p*<0.01. (**b**) Immunohistochemical staining for Ki-67 in HCC827 xenograft tumors in the different groups (scale bar, 100 μm). The percentage of Ki-67^+^ cells is presented as the mean ± SD of three sections. Ki-67^+^ cells were counted in 5 random fields per section. (**c**) The levels of methylated *P16* alleles (left chart) and *P16* expression (middle chart) in three representative tumors from each group, as assessed by MethyLight and qRT-PCR assays. (**d**) The levels of RB, phosphorylated RB (p-RB), and P16 proteins in three representative tumors from each group, as assessed by Western blot. Student’s t-test, \**p*<0.05, \*\**p*<0.01.

Moreover, the level of methylated *P16* alleles was significantly decreased in three representative P16-Dnmt tumors treated with palbociclib, while the level of *P16* mRNA was significantly increased in these tumors (Fig 4c). Changes in the *P16* mRNA level in control vector tumors could not be induced by palbociclib treatment. In addition, more P16 protein was detected in P16-Dnmt tumors treated with palbociclib than in those treated with control buffer, as evidenced by Western blot analysis (Fig 4d). Such a difference was not observed between control vector tumors with and without palbociclib treatment. In contrast, the p-RB levels in P16-Dnmt tumors treated with palbociclib were lower than those in the control vector tumors treated with palbociclib. Together, these results indicate that palbociclib can selectively inhibit the growth of *P16*-methylated cells *in vivo*.

## Discussion

On the basis of its good performance in clinical trials, the specific CDK4/6 inhibitor palbociclib has been approved by the FDA and EMA for HR-positive and HER2-negative advanced breast cancer. The P16 protein encoded by the *CDKN2A* gene is an endogenous CDK4/6 inhibitor. This gene is most frequently inactivated in human cancer genomes by copy number deletion, with a frequency of about 10%, and DNA methylation, with a frequency of approximately 30%. It has been reported that CDK4/6 inhibitors can effectively inhibit processes in tumor cells that lose endogenous inhibition of CDK4/6 because of *P16* gene deletion [14, 24]. However, it is not known whether *P16* inactivation by DNA methylation may similarly increase the sensitivity of cancer cells to these CDK4/6 inhibitors. In the present study, we found, for the first time, that the proliferation and growth of *P16*-methylated cancer cells can be selectively inhibited by palbociclib *in vitro* and *in vivo*.

Our bioinformatics analysis via mining public pharmacogenomic datasets indicates that the IC50 value of palbociclib is positively correlated with the transcription level of the *CDKN2A* gene in 522 cancer cell lines. As *P16* methylation can directly inactivate gene transcription [29], we found that similar to cell lines with *P16* deletion, *P16*-methylated cancer cell lines were more sensitive to palbociclib than *P16*-unmethylated cell lines. Notably, *P16*-specific methylation by P16-Dnmt directly increased the sensitivity of lung cancer cells to palbociclib *in vitro* and *in vivo*. Furthermore, we observed that palbociclib selectively inhibited the proliferation and growth of *P16*-methylated cells in P16-Dnmt tumors. Lower protein levels of p-RB were detected in P16-Dnmt tumors than in control vector tumors in nude mice treated with palbociclib. These phenomena strongly suggest that *P16*-methylated cells have increased sensitivity to CDK4/6 inhibitor drugs.

In the phase II clinical study PALOMA-1, it was found that patient selection based on *CCDN1* amplification and/or *P16* loss (*CDKN2A* deletion) did not improve outcomes of the administration of palbociclib plus letrozole in HR-positive breast cancer patients [37]. In the phase III PALOMA-2 trial, although the expression levels of genes in the Cyclin D-CDK4/6-RB pathway did not correlate with a benefit from palbociclib plus letrozole, 59 breast cancer patients without P16 expression exhibited a benefit [38, 39]. In another phase I study of the CDK4/6 inhibitor ribociclib (LEE011), *CCND1* amplification *vs*. *CDKN2A* and *CDKN2B* codeletion trended toward a longer vs. shorter treatment duration [12]. We recently reported that *P16* methylation could increase the resistance of lung (and stomach) cancers to paclitaxel treatment in a clinical trial [40]. In contrast, here, we reveal that *P16* methylation can increase the sensitivity of lung and stomach cancers to palbociclib. It is worth performing clinical trials to assess whether *P16* methylation could be used to predict the therapeutic efficacy of CDK4/6 inhibitors such as palbociclib in patients with lung and stomach cancers.

## Conclusion

This study demonstrates that *P16* methylation can increase the sensitivity of cancer cells to the CDK4/6 inhibitor palbociclib. *P16* methylation may be used as a predictive marker for sensitivity to CDK4/6 inhibitors.

## Acknowledgments

This work was financially supported by a grant from the Beijing Municipal Commission of Health and Family Planning (PXM2018_026279_00005) to DD.

## Author Contributions

**Conceptualization:** DD, PL.

**Data curation:** PL.

**Formal analysis:** PL.

**Funding acquisition:** DD.

**Investigation:** PL, JZ, LG, XZ.

**Methodology:** DD, PL.

**Project administration:** DD, PL.

**Resources:** DD.

**Supervision:** DD.

**Validation:** PL, XZ.

**Visualization:** PL, DD.

**Writing – original draft:** PL.

**Writing – review & editing:** DD, PL.

## Compliance with ethical standards

All animal experiments were approved by Peking University Cancer Hospital’s Institutional Animal Care and Use Committee and complied with the internationally recognized Animal Research: Reporting of *In Vivo* Experiments guidelines. This article does not contain any studies with human participants performed by any of the authors.

## List of abbreviations

7ZFP: seven zinc finger protein
BSA: bovine serum albumin
CCLE: Cancer Cell Line Encyclopedia
CDK: cyclin-dependent kinase
GDSC: Genomics of Drug Sensitivity in Cancer
HER2: human epidermal growth factor receptor 2
HR: hormone receptor
IC50: half-maximal inhibitory concentration
MSP: methylation-specific PCR
P16-Dnmt: P16-specific methyltransferase
PFS: progression-free survival
RB: retinoblastoma protein
RCN: relative copy number

## Supporting information

**S1 Fig. Selective inhibition of the proliferation of *P16*-methylated cancer cells by palbociclib *in vitro*.** The results of immunofluorescence and confocal microscopy analyses to directly detect alterations in the subpopulation of cells staining positive for P16 protein within HCC827 lung cancer cells stably transfected with P16-Dnmt. Scale bar, 400 μm. Student’s t-test, **p<0.01.

**S2 Fig. Selective inhibition of the proliferation of *P16*-methylated cancer cells by palbociclib *in vitro*.** The results of immunofluorescence and confocal microscopy analysis to directly detect alterations of the subpopulation of cells staining positive for P16 protein within the BGC823 gastric cancer cells stably transfected with P16-Dnmt. Scale bar, 400 μm. Student’s t-test, \*\**p*<0.01.

